# Optimization of anastomotic technique and gastric conduit perfusion with hyperspectral imaging in an experimental model for minimally invasive esophagectomy

**DOI:** 10.1101/2021.10.03.462901

**Authors:** F. Nickel, A. Studier-Fischer, B. Özdemir, J. Odenthal, L.R. Müller, S. Knödler, K.F. Kowalewski, I. Camplisson, M.M. Allers, M. Dietrich, K. Schmidt, G.A. Salg, H.G. Kenngott, A.T. Billeter, I. Gockel, C. Sagiv, O.E. Hadar, J. Gildenblat, L. Ayala, S. Seidlitz, L. Maier-Hein, B.P. Müller-Stich

**Affiliations:** Department of General, Visceral, and Transplantation Surgery, Heidelberg University Hospital, Heidelberg, Germany; HIDSS4Health - Helmholtz Information and Data Science School for Health, Heidelberg and Karlsruhe, Germany; School of Medicine, Heidelberg University, Heidelberg, Germany; Division of Computer Assisted Medical Interventions, German Cancer Research Center (DKFZ), Heidelberg, Germany; Faculty of Mathematics and Computer Science, Heidelberg University, Heidelberg, Germany; Department of Urology, Medical Faculty of Mannheim at the University of Heidelberg, Mannheim, Germany; Division of Biology and Biological Engineering, California Institute of Technology, Pasadena, USA; Department of Anaesthesiology, Heidelberg University Hospital, Heidelberg, Germany; Department of Anaesthesiology and Intensive Care Medicine, Essen University Hospital, Essen, Germany; Department of Visceral, Transplantation, Thoracic and Vascular Surgery, Leipzig University Hospital, Leipzig, Germany; DeePathology Ltd., Ra’anana, Israel; Medical Faculty, Heidelberg University, Heidelberg, Germany

**Author notes:** **Correspondence to:** Prof. Dr. Beat P. Mueller-Stich, Department of General, Visceral, and Transplantation Surgery, Heidelberg University Hospital, Im Neuenheimer Feld 420, 69120 Heidelberg, Germany. These authors contributed equally (F. Nickel, A. Studier-Fischer).

**Keywords:** hyperspectral imaging, minimally invasive esophagectomy, anastomotic insufficiency, gastric conduit, linear stapling, linear stapled anastomosis, esophagogastric anastomosis, translational research, porcine model, tissue perfusion

## Abstract

**Objective:** To optimize anastomotic technique and gastric conduit perfusion with hyperspectral imaging (HSI) for total minimally invasive esophagectomy (MIE) with linear stapled anastomosis.

**Summary Background Data:** Esophagectomy is the mainstay of esophageal cancer treatment but anastomotic insufficiency related morbidity and mortality remain challenging for patient outcome.

**Methods:** A live porcine model (n=50) for MIE was used with gastric conduit formation and linear stapled side-to-side esophagogastrostomy. Four main experimental groups differed in stapling length (3 vs. 6 cm) and anastomotic position on the conduit (cranial vs. caudal). Tissue oxygenation around the anastomotic site was evaluated using HSI and was validated with histopathology.

**Results:** The tissue oxygenation (ΔStO_2_) after the anastomosis remained constant only for the short stapler in caudal position (−0.4± 4.4%, n.s.) while it dropped markedly in the other groups (short-cranial: -15.6± 11.5%, p=0.0002; long-cranial: -20.4± 7.6%, p=0.0126; long-caudal: -16.1± 9.4%, p<0.0001) Tissue samples from deoxygenated stomach as measured by HSI showed correspondent eosinophilic pre-necrotic changes in 35.7± 9.7% of the surface area.

**Conclusions:** Tissue oxygenation at the anastomotic site of the gastric conduit during MIE is influenced by stapling technique. Optimal oxygenation was achieved with a short stapler (3 cm) and sufficient distance of the anastomosis to the cranial end of the gastric conduit. HSI tissue deoxygenation corresponded to histopathologic necrotic tissue changes. These findings allow for optimization of gastric conduit perfusion and anastomotic technique in MIE.

**Level of Evidence:** Not applicable. Translational animal science. Original article.

## 1 Introduction

Treatment for esophageal cancer is stage-dependent and can include endoscopic resection, esophagectomy and chemotherapy as well as radiation. The mainstay of treatment for any resectable esophageal cancer is esophagectomy^1^. Technical and technological improvements such as minimally invasive surgery and stapled anastomosis have led to reduced morbidity and mortality of esophagectomy over the last decades. There is sufficient evidence nowadays showing that oncological outcomes between open esophagectomy (OE) and minimally invasive technique are at least equivalent.^2-4^ Minimally invasive esophagectomy (MIE) has especially shown improved pulmonary complication rates.^5,6^ Currently, there is no official gold standard technique for the creation of intrathoracic esophagogastric anastomosis after Ivor-Lewis esophagectomy and the choice of technique depends on the localization of the tumor as well as the surgeon’s preference and experience.^7^ The perfusion and tissue oxygenation of the gastric conduit and integrity of the anastomosis are critical factors for short- and long-term outcome.

While the end of the gastric conduit can more often experience perfusion-associated problems, perfusion is usually sufficient at the esophageal stump. For the linear-stapled technique in MIE the anastomosis is constructed in a side-to-side fashion. Besides popularity of end-to-side circular stapled anastomosis for OE and robotic-assisted MIE, the linear stapled anastomosis provides a method with good feasibility and potential for standardization in conventional MIE. The linear stapled technique is well established in laparoscopic bariatric surgery and can serve as a mainstay technique for conventional MIE, especially if a robotic system is not available. There is currently no evidence for the exact placement of the linear stapled anastomosis during MIE, although this may influence tissue perfusion and oxygenation and therefore the risk of complications.

Hyperspectral Imaging (HSI) is a novel imaging technique that can estimate tissue oxygenation and microcirculatory perfusion by measuring reflectance intensity separately at different wavelengths^8^ (**Figure 1**). It then calculates color-coded index images that can be used to measure tissue oxygenation and perfusion and enables microvascular evaluation of organ perfusion^9-22^, tissue identification^11,23-26^ and assessment of different tissue conditions^25,27^. It is therefore well suited to evaluate different technical aspects of esophagectomy such as anastomotic stapling techniques regarding tissue perfusion as risk factors for anastomotic leakage.

**Figure 1.**
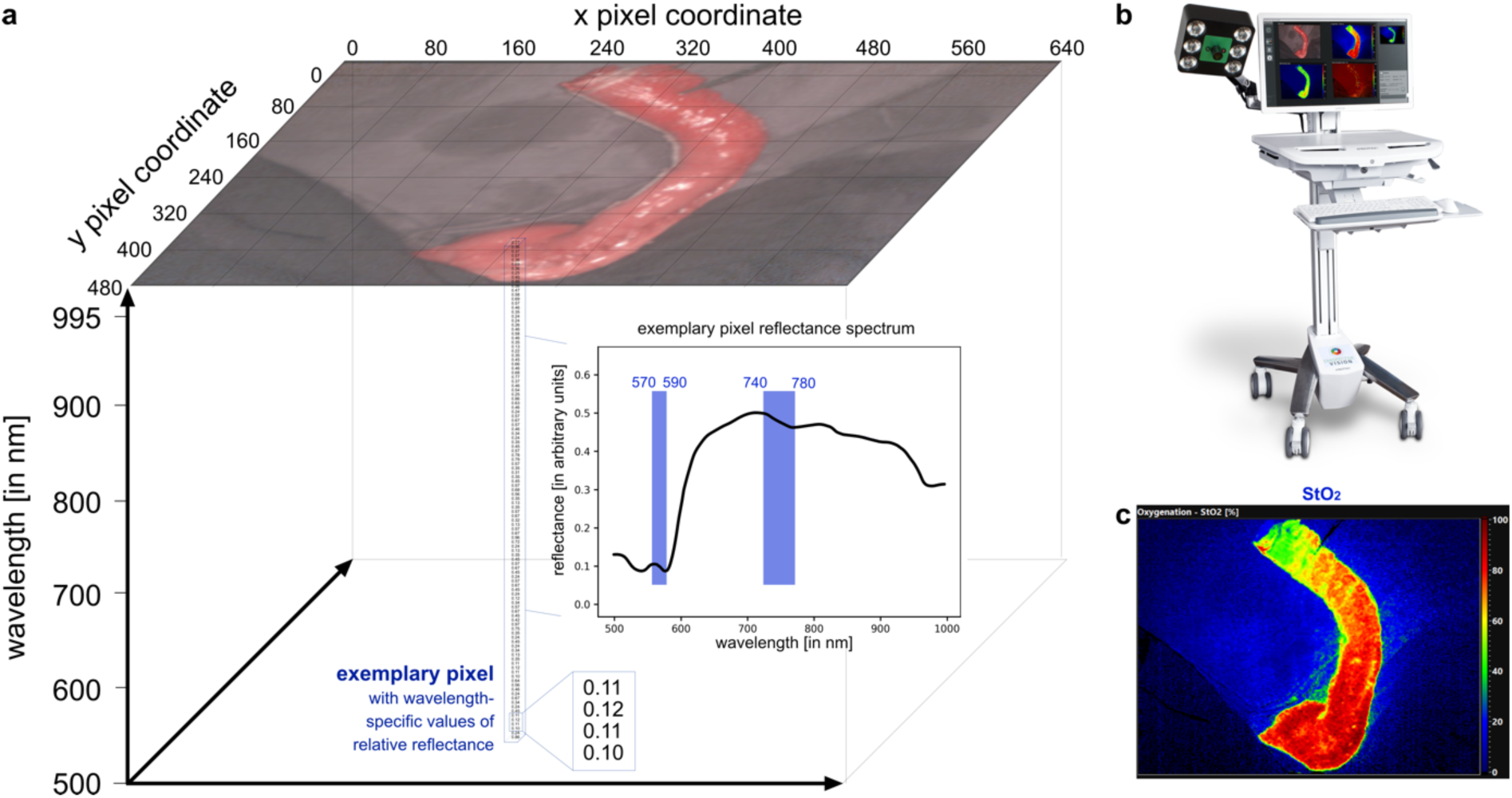
Hyperspectral data cube and camera system. **a**, visualization of a three-dimensional hyperspectral datacube indicating the spectral bands relevant for the calculation of the hyperspectral oxygenation index (StO_2_). **b**, TIVITA® Tissue hyperspectral camera system with the kind permission from Diaspective Vision GmbH. **c**, color-coded picture of StO_2_ index.

The aim of the present study was to assess different stapling positions and techniques and their effect on tissue oxygenation in a porcine model for MIE. The optimization of gastric conduit perfusion and anastomotic technique in an experimental model for MIE will serve to improve understanding of gastric conduit physiology and to reduce complications for patients requiring esophagectomy.

## 2 Materials and methods

### 2.1 Porcine model and surgical procedure

The experiments were approved by the Committee on Animal Experimentation of the regional council Baden-Württemberg in Karlsruhe (G-161/18 and G-262/19). All pigs were managed according to German laws for animal use and care and according to the directives of the European Community Council (2010/63/EU) and ARRIVE guidelines.^28^

In order to obtain a first impression regarding spectral differences between physiological (A=39; n=849) and deoxygenated (A=15; n=117) stomach, hyperspectral recordings were taken before and after blood supply was completely dissected (A=50; n=966 in total). A always indicates the number of animals; n always indicates the number of independent measurements in total (**Figure 2**).

**Figure 2.**
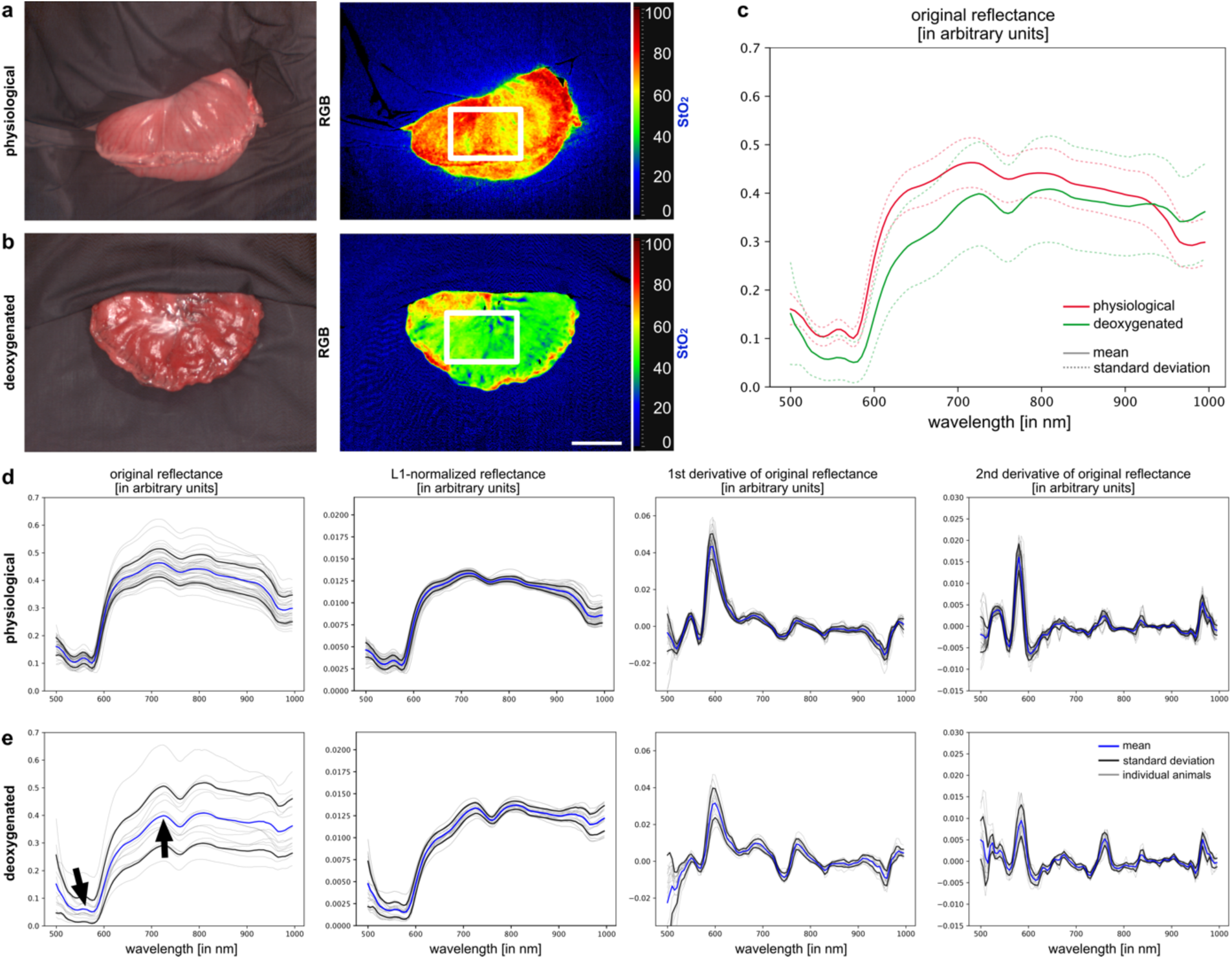
Hyperspectral imaging characterization of physiological and deoxygenated stomach. TIVITA® color-coded index images for oxygenation (StO_2_) and relative reflectance intensities across wavelengths. **a**, images for physiological stomach (A=39; n=849). **b**, images for deoxygenated stomach (A=15; n=117). **c**, mean spectra in comparison. **d**, reflectance and derivatives of physiological stomach. **e**, reflectance and derivatives of deoxygenated stomach. Black arrows indicate the spectral areas that are mainly influenced by the oxygenation status of hemoglobin. Mean and standard deviation are calculated across individual animals. A always indicates the number of animals; n always indicates the number of independent measurements in total. Scale bar always equals 5 cm if not stated otherwise.

For the purpose of this project’s main research question, the specific interest was the anastomotic site at the end of the gastric conduit as this is known to be the critical location regarding perfusion, and the anastomotic site at the esophageal stump was not of interest. Consequently, only the abdominal procedure of Ivor-Lewis esophagectomy was performed in an open fashion via midline laparotomy, but with the technical aspects as used in MIE, i.e. linear stapled anastomosis. A standardized gastric conduit was created with stapling devices. For gastric conduit perfusion, the only remaining vessels of supply were the right gastroepiploic artery and vein. During all of the following steps, hyperspectral images of the conduits were recorded.

In order to impair blood supply of the conduit, strong magnets with the same dimensions as typical stapling devices were applied in different ways. While one magnet was placed intraluminally, the counterside magnet was placed on the ventral surface of the gastric conduit. In pilot studies, these magnets were shown to be strong enough to completely inhibit tissue blood flow and therefore also adequately simulate the anastomotic formation with stapling devices (**Supplementary Text 3** and **Supplementary Figure 3**). There were magnets in two lengths, namely 3 cm and 6 cm, in order to simulate 3 cm and 6 cm linear stapled anastomoses.

In theory, there are two conceivable ways of perfusion that can contribute to the blood supply of the gastric conduit. These are longitudinal tissue capillary perfusion on the one hand (**Figure 3a**), and trans-arterial and subsequent transverse tissue capillary perfusion via the gastroepiploic artery on the other (**Figure 3b**). In order to investigate the primary way of blood supply, gastric conduit perfusion was impaired intentionally with transverse and longitudinal magnets. In a separate experiment, the gastroepiploic artery was clamped at different levels and subsequently released to investigate effects of impaired right gastroepiploic artery perfusion (**Figure 3c**).

**Figure 3.**
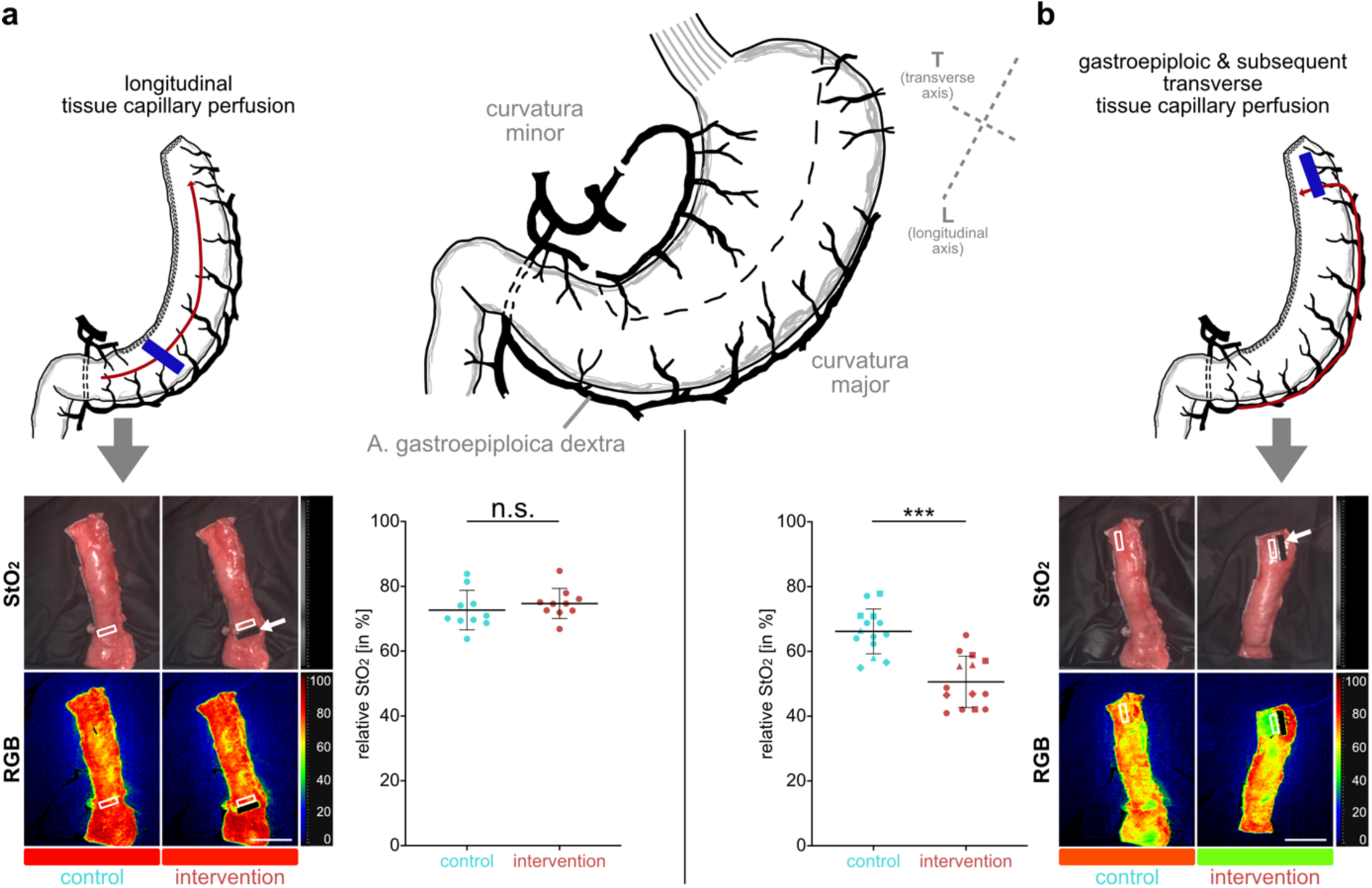
Evaluation of possible ways of gastric conduit perfusion with hyperspectral imaging. **a**, longitudinal tissue capillary perfusion relative to the conduit (A=10; n=10) and schematic depiction of relevant gastric anatomy with longitudinal (L) and transverse (T) axis of the later conduit. **b**, trans-arterial and subsequent transverse tissue capillary perfusion relative to the conduit via the gastroepiploic artery (A=10; n=14). Arrows indicate the site of perfusion inhibition through magnets. White boxes indicate regions of interest (ROIs) for measurements. A paired t-test was applied; n.s. is not significant, *** is p≤0.001. Numbers in the box plots were obtained from the ROIs. Different animals are coded in different shapes; circles are different animals. Graphs depict mean and standard deviation.

To investigate the four main experimental groups of this work, the linear incision on the ventral side of the gastric conduit usually caused by the linear stapling device in MIE was now simulated with magnets that had the same dimensions as typical stapling devices (**Figure 4**).

**Figure 4.**
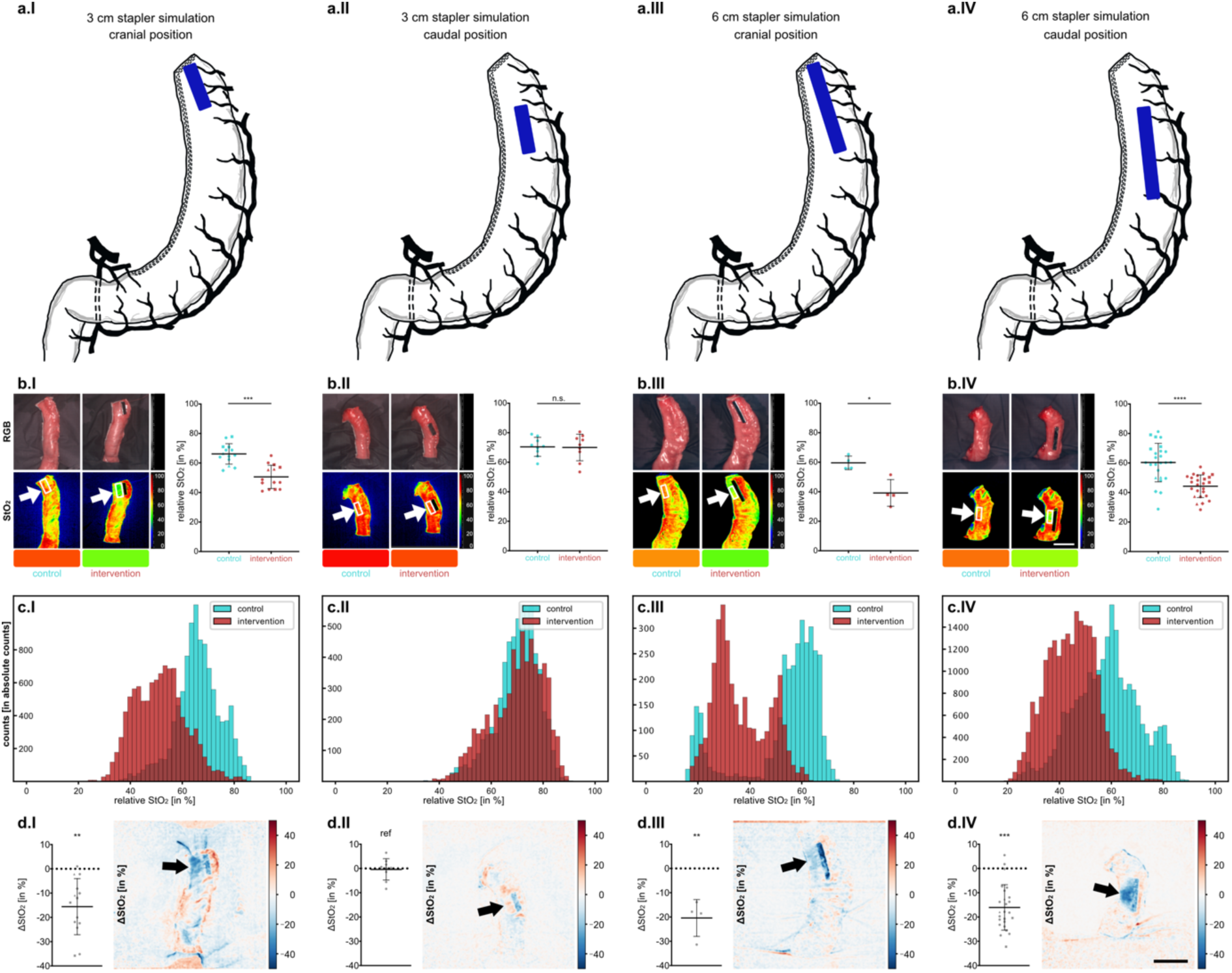
Effects of linear stapled esophagogastric anastomotic stapling position and length. **I**, short (3 cm) cranial stapler simulation (A=10; n=14). **II**: short (3 cm) caudal stapler simulation (A=9; n=9). **III**: long (6 cm) cranial stapler simulation (A=3; n=4). **IV**: long (6 cm) caudal stapler simulation (A=20; n=25). **a**, schematic drawing. **b**, exemplary visualization of oxygenation and corresponding quantification. White boxes and arrows indicate regions of interest (ROIs) for measurements. For **b** a paired t-test was performed. **c**, histograms of all StO_2_ values without hierarchical structure. **d**, quantification of the StO_2_ difference in the ROIs between control and intervention and exemplary color-coded visualization. Black arrows indicate the ROI next to the magnet. For **d** a one-way ANOVA was performed; only significant differences to group II (short caudal) as reference were highlighted; all other comparisons were not significant; ref indicates reference group for statistical testing. * is p≤0.05, ** is p≤0.01, *** is p≤0.001, **** is p≤0.0001, n.s. is not significant. Numbers in the box plots were obtained from the ROIs. Different animals are coded in different shapes; circles are different animals. Graphs depict mean and standard deviation.

The four main experimental groups that represent possible stapling methods for the linear stapled anastomosis were (group I) short stapler (3 cm length) with cranial stapling position, (group II) short stapler with caudal stapling position, (group III) long stapler (6 cm length) with cranial stapling position, and (group IV) long stapler with caudal stapling position (**Figure 4a**). While the cranial stapling position describes the closest possible position of the intraluminal magnet to the cranial staple line of the gastric conduit with no well perfused tissue remaining at the end of the conduit, the caudal stapling position describes the magnet position 3 cm below, so that there is 3 cm of well perfused gastric conduit tissue still above. HSI recordings were done for the physiological gastric conduit and then with the magnets in place to simulate the anastomosis after 2 minutes. For a subgroup analysis the gastric conduit width was performed with a standardized diameter of 2 cm for the thin conduit group and 4 cm for the wide conduit group (**Supplementary Figure 3**).

### 2.2 Hyperspectral imaging and statistical analysis

HSI data was acquired with the TIVITA® Tissue system (Diaspective Vision GmbH). It provides a spectral resolution of 5 nm in the range from 500 nm to 995 nm **(Figure 1)**. Six integrated halogen lamps provide a standardized illumination. Numerical values stored in the datacube represent reflectance values at every single pixel for every wavelength in arbitrary units. The computed tissue parameter index images include the hyperspectral oxygenation index (StO_2_) and underlying formulas can be reviewed in cited literature^8^ **(Figure 1)**. For visualization purposes, the control StO_2_ images were subtracted from the intervention StO_2_ images as necessary, resulting in ΔStO_2_ images.

Mean and standard deviation (SD) reflectance spectra of physiological and deoxygenated stomach (**Figure 2**) across animals were obtained by calculating the median spectrum over every pixel within the annotated region of interest (ROI) of one image and then the mean spectrum over every image within one pig. For the comparison of gastric conduit recordings before (control) and after magnet application for anastomosis simulation (intervention) both were registered using the Optical flow algorithm based on the annotation of the contour of each conduit ^29-31^ (**Supplementary Text 1** and **Supplementary Figure 1**), a ROI of identical size of 16×50 pixels for each recording in the intervention was manually placed and the ROI in the control was set automatically according to the registration. For any figures reporting StO_2_, the mean ± SD over every pixel within the ROI of one image was calculated and then the mean over every image of one experimental group. For the analysis of the four main experimental groups, a ROI for each recording in the intervention group was defined left to the magnet. Baseline measurements were taken from respective registered control recordings without magnet.

All displayed StO_2_ values were obtained as relative values from the official formula used by the HSI camera as cited in the literature^8^. Different animals were coded in different shapes; in analyses of hyperspectral data circles are measurements from different animals. A p-value ≤0.05 was considered statistically significant. In case of parametric data, paired and unpaired t-test was used. In case of non-parametric data, Mann-Whitney test was used for unpaired and Wilcoxon matched-pairs signed rank test for paired data. For comparisons of multiple groups, one-way ANOVA was used in case of parametric normal distribution while Kruskal-Wallis was used in case of non-parametric distribution. p-values were adjusted for multiple testing.

### 2.3 Pathohistological analysis

In order to evaluate whether the differences in HSI oxygenation have any biological relevance and reflect histopathological changes, tissue in the main experimental group IV was sampled from physiological stomach (R1) and from 3 distinctive regions after 6 hours of magnet-induced ischemia (R2-4) (**Figure 5**). R2 serves as baseline of physiological tissue 6 hours into the surgery. R4 is defined by lowest HSI oxygenation alongside the magnet. R3 is the intermediate region. All tissue samples were formalin-fixated and hematoxylin-eosin stained. Slides were scanned using widefield microscopy (Zeiss) and the amount of mucosal eosinophilic pre-necrotic areas was objectified with the specialized software DeePathology™ STUDIO (Deepathology Ltd., Israel) (**Figure 5**). Description of methods for vascular corrosion casting and scanning electron microscopy (SEM) for further pathomechanistic investigation can be found in the supplement **(Supplementary Text 4** and **Supplementary Figure 4)**.

**Figure 5.**
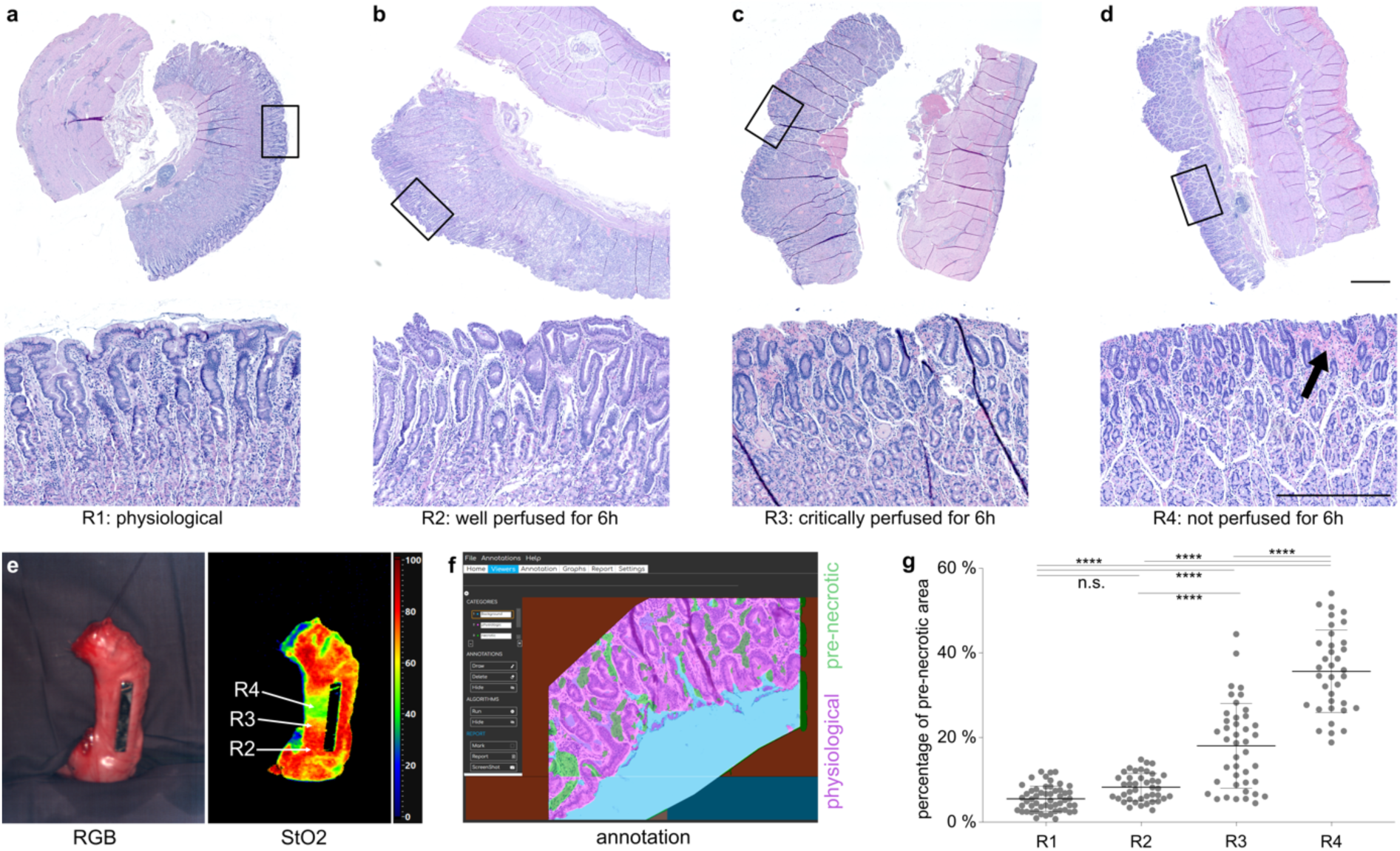
Pathohistological samples in correlation to hyperspectral findings. **a-d**, Pathohistological HE samples showing whole slides and representative mucosal areas for R1: physiological (A=4; n=54), R2: well perfused for 6h (A=4; n=41), R3: critically perfused for 6h (A=4; n=42) and R4: not perfused for 6h (A=4; n=36). **e**, hyperspectral recording of gastric conduit with impaired perfusion due to magnet-induced ischemia. **f**, DeePathology™ STUDIO with visualization of physiological areas in magenta and eosinophilic pre-necrotic areas in green. **g**, quantification of eosinophilic pre-necrotic areas in percentage of surface area; different animals are coded in different shapes. Graph depicts mean and standard deviation. Ordinary one-way ANOVA with multiple comparisons was used. n.s. is not significant, **** is p≤0.0001. Upper scale bar indicates 1 mm, lower scale bar indicates 500 μ

## 3 Results

### 3.1 Physiological and deoxygenated stomach

Differences between physiological stomach and deoxygenated stomach could be clearly detected in the HSI StO_2_ images (**Figure 2**). Distinct changes in the reflectance spectrum were seen between 550 and 600 nm and above 700 nm.

### 3.2 Capillary and gastroepiploic perfusion

Inhibiting longitudinal tissue capillary perfusion by using a transverse magnet position (**Figure 3a**) did not yield any significant difference between the StO_2_ values of control (72.7± 6.1%) and intervention (74.8± 4.6%). However, when limiting transarterial and subsequent transverse tissue capillary perfusion via the gastroepiploic artery with a longitudinal magnet (**Figure 3b**), StO_2_ levels significantly dropped from 66.2± 6.9% to 50.6± 7.9% (p=0.0002).

### 3.3 Different stapler sizes and positions influence tissue oxygenation of linear stapled anastomosis during esophagectomy

Due to results of aforementioned experiments regarding capillary and gastroepiploic perfusion, the region with the lowest StO_2_ values on the gastric conduit during intervention can be expected on the left side of the magnet downstream to the perfusion coming transversely from the gastroepiploic artery. The four main experimental groups representing possible stapling methods for constructing the anastomosis during MIE were evaluated for changes in tissue oxygenation (**Figure 4**). Oxygenation levels were significantly reduced in group I (short-cranial) from 66.2± 6.9% to 50.6± 7.9% (ΔStO_2_ = - 15.6± 11.5%, p=0.0002), in group III (long-cranial) from 59.6± 4.5% to 39.3± 9.0% (ΔStO_2_ = -20.4± 7.6%, p=0.0126) and in group IV (long-caudal) from 60.3± 13.0% to 44.3± 7.7% (ΔStO_2_ = -16.1± 9.4%, p<0.0001) (**Figure 4b.I, III and IV**). Only in group II (short-caudal) changes from 70.5± 6.4% to 70.1± 9.0% were not significant (ΔStO_2_ = -0.4± 4.4%, n.s.) (**Figure 4b.II**). ΔStO_2_ images (**Figure 4d**) were all significantly different from group II, which served as a reference with values of -0.4± 4.4%. ΔStO_2_ values were -15.6± 11.5% (p=0.0021) for group I, -20.4± 7.6% (p=0.0043) for group III and - 16.1± 9.4% (p=0.0004) for group IV. All other comparisons were not significantly different. This allows for the conclusion that only for a short stapler with caudal position, tissue oxygenation does not significantly drop after tissue dissection.

### 3.4 Pathohistological correlation to hyperspectral findings

The area of pre-necrotic changes in the samples were quantified with 5.5± 3.0% for R1, 8.3± 3.2% for R2, 18.1± 10.0% for R3 and 35.7± 9.7% for R4. All differences were significant as indicated (p<0.0001).

## 4 Discussion

Anastomotic integrity is the Achilles’ heel of esophagectomy and other gastrointestinal procedures. It is multifactorial with the most important influencing factors being sufficient tissue perfusion, lack of tension, asepsis and quality of surgical sutures and stitches ^32^. The present experimental study investigated several technical aspects of linear stapled esophagogastric anastomosis used in Ivor Lewis MIE focusing on anastomotic perfusion. Linear stapled side-to-side esophagogastric anastomosis is a viable alternative to open or robotic circular stapled technique and especially valuable if advantages of MIE are desired despite the lack of a robotic system ^33^. In bariatric surgery the linear stapled side-to-side anastomosis has proven to be a reproducible and standardizable technique with good clinical performance causing it to be the focus of this investigation with regard to MIE ^34,35^. The present study established a translational method and blueprint for the systematic evaluation of different technical aspects of the gastric conduit and anastomosis for esophagectomy in a porcine model. HSI was used as a tool for objective tissue perfusion evaluation intended for optimization of surgical technique.

In order to provide a first impression of spectral differences between physiological and non-perfused gastric tissue, HSI was utilized to record both of those conditions. Spectral reflectance was clearly different with a double-peak between 550 and 590 nm and higher values at 680 nm for physiological tissue compared to non-perfused gastric tissue as seen in **Figure 2**. These findings are in line with the descriptions of optical properties of oxygenated and deoxygenated hemoglobin in current literature ^36-38^. This initial analysis serves as a reference point for the subsequent experiments as spectral reflectance of the critically perfused areas on the gastric conduit next to the stapling site is expected to be similar compared to the spectral reflectance of non-perfused gastric tissue.

As a first step, basic principles of gastric conduit perfusion dynamics were explored by impairing the two possible ways of blood supply i.e. longitudinal capillary perfusion in the gastric conduit tissue along the greater curvature or alternatively perfusion along the gastroepiploic artery and subsequently capillaries transversely to the conduit axis as illustrated in **Figure 3**. It could be shown that there is no sign of relevant longitudinal capillary perfusion, but proof of perfusion along the gastroepiploic artery as seen in **Figure 3, Supplementary Text 2** and **Supplementary Figure 2**. This observation matches results from human anatomical studies, which showed that larger capillaries are distributed perpendicular to both gastric curvature sides and only small capillary branches parallel to both curvature sides originate from these to form a vascular network throughout the stomach wall ^39^. The dominant direction of flow is therefore not a random pattern along the longitudinal axis of the conduit. Thus, perfusion mainly occurs through the right gastroepiploic artery and subsequent capillaries along the transverse axis of the conduit.

Especially the distal end of the conduit also relies on a sufficient collateral connection of right and left gastroepiploic artery. Impairment in this collateral anatomy might cause insufficient perfusion of the conduit’s end with the consequences of necessary resection and shortening possibly leading to anastomotic tension. A human study found the right gastroepiploic artery to be more dominant in that the mean cross-sectional area was found to be 3.31±1.71 mm^2^ compared to 1.33±1.01 mm^2^ for the left gastroepiploic artery and 0.51±0.28 mm^2^ for the collateral connection between both ^40^. Similar observations were made during the present porcine study in the sense that the right gastroepiploic artery was the dominant vessel supplying the gastric conduit. Furthermore, different human anatomical studies report different rates of actual collateral anastomotic connections between right and left gastroepiploic artery reaching from as high as 95% with an arterial arch (70%) or mesh-like anastomosis (25%) ^41^ over 70% ^42^ down to as low as only 60% ^43^. The knowledge of these anatomical key points is crucial for gastric conduit geometry and formation. It can therefore be concluded that in the porcine model for esophagectomy with reconstruction via a gastric conduit, the relevant amount of perfusion is almost exclusively due to the right gastroepiploic artery and its transverse capillary branches. The possible length of the gastric conduit therefore also depends on size and collateral connection between both gastroepiploic arteries.

An in-depth analysis of different anastomotic linear stapler techniques showed that the most promising stapling technique in order to preserve the optimal tissue perfusion measured with HSI is a short stapler (3 cm as compared to 6 cm) with sufficient distance (3 cm) of the cranial end of the anastomosis to the stapled end of the conduit as depicted in **Figure 4**. The exact distances and sizes of staple lines are not always reported in clinical practice and no publications could be found addressing the proximity of the anastomosis towards the cranial end of the conduit. However, there were two comparable clinical trials investigating MIE Ivor Lewis esophagectomy of which one used a 6 cm linear stapler ^44^, while the other used a 3.5 cm linear stapler ^45^. Within the 124 patients with a 6 cm anastomosis 9 anastomotic leaks occurred (7.3%), while within the 104 patients with a 3 cm anastomosis only 4 patients (3.8%) experienced anastomotic leakage. Overall postoperative complications were similar with 51.6% compared to 50.0%. Although one could expect higher stricture rates for a smaller stapler, the 3 cm anastomosis only had 1 stricture (<1.0%) compared to the 6 cm anastomosis with 6 patients (5.1%) with anastomotic stricture requiring endoscopic intervention. This is in line with the current study indicating that hypoperfusion of the perianastomotic tissue is more prevalent with longer staplers used for anastomosis, which can lead to anastomotic insufficiency on the short term, but also to scarring and strictures on the long term. It therefore seems that patients with a shorter linear stapler anastomosis benefit from improved anastomotic perfusion with potentially lower risk of anastomotic leakage while not bearing additional risks for stricture.

In order to show biological relevance and potential for clinical translation of the HSI-based StO_2_ measurements, the effect on tissue necrosis was investigated by histopathological examination in the present study. Differences between gastric conduit groups in histologically-observed tissue reactions to induced tissue ischemia clearly corresponded to the observed different hyperspectral StO_2_ values. It could be shown that lower StO_2_ values in HSI corresponded to higher percentages of pre-necrotic stains in the gastric mucosa of histopathological samples, hereby indicating significant tissue damage through insufficient perfusion (**Figure 5**). Physiologically high StO_2_ values corresponded to physiologically low levels of pre-necrotic tissue changes. A typical drop in StO_2_ from physiological HSI values between 60% and 70% as described in other studies ^46^ to below 45% during magnet-induced ischemia resulted in a pre-necrotic demarcation of gastric tissue of 35.7± 9.7% of histological surface area. This tendency also applied incrementally for the stages in between, so that with lower StO_2_ values between 45% and 60%, there were correspondingly greater surface areas between 10% and 35% with pre-necrotic changes. Histopathological analysis as seen in **Figure 5** therefore confirmed the relevance of HSI StO_2_ changes for assessment of real tissue condition.

There are several limitations of this study. First, all of the experiments were performed in a porcine model and although the porcine anatomy is very similar to the human anatomy, clinical translation of these results has to be proven. Secondly, for patients receiving esophagectomy there are additional factors that influence anastomotic integrity. While ideal stapling position and gastric conduit formation as described in this work may represent one decisive factor for anastomotic healing, other factors also come into play such as a tension-free anastomosis, fluid volume management, vasopressors, blood pressure, respiratory function and resulting oxygen supply, comorbidities and nutrition amongst others. It is therefore essential for the successful treatment of patients to take all the relevant factors into consideration in order to obtain the best possible outcome. Explicitly for the correlation of HSI StO_2_ values and histopathological tissue changes, it is important to mention that this type of validation will have to be repeated for different hyperspectral index picture values, organs and even species. These limitations are essential, but when accounted for, the present animal model for MIE can be used as a blueprint for the systematic investigation of gastric conduit perfusion dynamics for multiple research questions yet to come.

## 5 Conclusion

A porcine model for optimization of gastric conduit formation with analysis of tissue oxygenation and microcirculation using the novel intraoperative imaging technique of HSI was developed. Different aspects of anastomotic linear stapling technique for minimally invasive Ivor Lewis esophagectomy were evaluated. A shorter linear stapler (3 cm compared to 6 cm) with at least 3 cm distance of the anastomosis to the cranial end of the gastric conduit provided optimal results in terms of tissue oxygenation and perfusion and is thus the most promising technique in terms of anastomotic healing. This animal model can be used as a blueprint for further systematic investigation of gastric conduit perfusion and optimization of anastomotic technique. Clinical trials are essential to investigate translation to optimized clinical outcome with reduction of anastomotic insufficiency and conduit necrosis.

## Abbreviations

HSI: hyperspectral imaging
i.v.: intravenous
MIE: minimally invasive esophagectomy
OE: open esophagectomy
ROI: region of interest
SD: standard deviation
SEM: scanning electron microscopy
StO_2_: hyperspectral oxygenation index

## Acknowledgements

The authors gratefully acknowledge the data storage service SDS@hd supported by the Ministry of Science, Research and the Arts Baden-Württemberg (MWK) and the German Research Foundation (DFG) through grant INST 35/1314-1 FUGG and INST 35/1503-1 FUGG. Furthermore, the authors gratefully acknowledge the support from the Willy Robert Pitzer Foundation and the Heidelberg Foundation of Surgery.

## Disclosure Information

Research funding: There was financial support from the Willy Robert Pitzer Foundation and the Heidelberg Foundation of Surgery for this project. Conflict of interest: Authors state no conflict of interest. Felix Nickel reports support for courses and travel from Johnson and Johnson, Medtronic, Intuitive Surgical, Cambridge Medical Robotics and KARL STORZ as well as consultancy fees from KARL STORZ. We acknowledge support from Medtronic for providing the stapling devices. Published data will be made available upon reasonable request to the corresponding author.

## Author contribution

ASF, FN and BPM had the original idea for the project. ASF and FN developed the project. ASF coordinated the whole project. ASF, FN, GS and KS performed the initial review of existing literature and the planning. ASF, KFK, BÖ, BPM and FN performed the surgeries. ASF and GS did the casting and the SEM recordings. ASF, IC, BÖ and JO developed the Python codes for data structure and annotation. ASF, SK, MMA, MD and KS annotated data. Image processing was performed by LRM, ASF, SK, SS, LA and LMH. ASF, MMA, GS, BÖ, JO, SK, KS, SS, LA, LRM and LMH analyzed and interpreted data. Statistical analysis was performed by ASF, KFK and FN. CS, OEH and JG supported the pathohistological analysis. ASF and FN drafted the figures and wrote the manuscript. SS, LRM, BPM, HK, IG, AB, CS, OEH and LMH revised the manuscript. All authors have read and approved the final manuscript.

## Registration of research studies

Not applicable.

## Notes

### Competing Interest Statement

The authors have declared no competing interest.

## References

1 Siegel, R. L., Miller, K. D. & Jemal, A. Cancer statistics, 2020. CA: A Cancer Journal for Clinicians 70, 7–30, doi:https://doi.org/10.3322/caac.21590 (2020).

2 Biebl, M., Andreou, A., Chopra, S., Denecke, C. & Pratschke, J. Upper Gastrointestinal Surgery: Robotic Surgery versus Laparoscopic Procedures for Esophageal Malignancy. Visc Med 34, 10–15, doi:10.1159/000487011 (2018).

3 Monig, S. et al. Early esophageal cancer: the significance of surgery, endoscopy, and chemoradiation. Annals of the New York Academy of Sciences, doi:10.1111/nyas.13955 (2018).

4 Müller-Stich, B. P. et al. Meta-analysis of randomized controlled trials and individual patient data comparing minimally invasive with open oesophagectomy for cancer. British Journal of Surgery, doi:10.1093/bjs/znab278 (2021).

5 Yibulayin, W., Abulizi, S., Lv, H. & Sun, W. Minimally invasive oesophagectomy versus open esophagectomy for resectable esophageal cancer: a meta-analysis. World J Surg Oncol 14, 304, doi:10.1186/s12957-016-1062-7 (2016).

6 van der Sluis, P. C. et al. Robot-assisted Minimally Invasive Thoracolaparoscopic Esophagectomy Versus Open Transthoracic Esophagectomy for Resectable Esophageal Cancer: A Randomized Controlled Trial. Ann Surg 269, 621–630, doi:10.1097/sla.0000000000003031 (2019).

7 Taurchini, M. & Cuttitta, A. Minimally invasive and robotic esophagectomy: state of the art. J Vis Surg 3, 125, doi:10.21037/jovs.2017.08.23 (2017).

8 Holmer, A., Marotz, J., Wahl, P., Dau, M. & Kammerer, P. W. Hyperspectral imaging in perfusion and wound diagnostics - methods and algorithms for the determination of tissue parameters. Biomed Tech (Berl), doi:10.1515/bmt-2017-0155 (2018).

9 Grambow, E. et al. Evaluation of peripheral artery disease with the TIVITA(R) Tissue hyperspectral imaging camera system. Clinical hemorheology and microcirculation 73, 3–17, doi:10.3233/ch-199215 (2019).

10 Jansen-Winkeln, B. et al. [Hyperspectral imaging of gastrointestinal anastomoses]. Chirurg 89, 717–725, doi:10.1007/s00104-018-0633-2 (2018).

11 Köhler, H., Jansen-Winkeln, B., Chalopin, C. & Gockel, I. Hyperspectral imaging as a new optical method for the measurement of gastric conduit perfusion. Diseases of the Esophagus 32, 1–1, doi:10.1093/dote/doz046 (2019).

12 Langner, I. et al. Hyperspektralimaging demonstriert mikrozirkulatorische Effekte postoperativer Ergotherapie bei Patienten mit Morbus Dupuytren. Handchirurgie, Mikrochirurgie, plastische Chirurgie : Organ der Deutschsprachigen Arbeitsgemeinschaft fur Handchirurgie : Organ der Deutschsprachigen Arbeitsgemeinschaft fur Mikrochirurgie der Peripheren Nerven und Gefasse 51, 171–176, doi:10.1055/a-0916-8635 (2019).

13 Marotz, J. et al. Extended Perfusion Parameter Estimation from Hyperspectral Imaging Data for Bedside Diagnostic in Medicine. Molecules 24, 4164, doi:10.3390/molecules24224164 (2019).

14 Mehdorn, M. et al. Hyperspektrales Imaging zur Diskrimination des Resektionsausmaßes im Rahmen der akuten Mesenterialischämie: Eine Fallserie. Z Gastroenterol 57, KV 82, doi:10.1055/s-0039-1695182 (2019).

15 Sicher, C. et al. Hyperspectral imaging as a possible tool for visualization of changes in hemoglobin oxygenation in patients with deficient hemodynamics – proof of concept. Biomedical Engineering / Biomedizinische Technik 63, 609, doi:https://doi.org/10.1515/bmt-2017-0084 (2018).

16 Sucher, R. et al. Hyperspectral Imaging (HSI) in anatomic left liver resection. International journal of surgery case reports 62, 108–111, doi:https://doi.org/10.1016/j.ijscr.2019.08.025 (2019).

17 Wild, T. et al. Hyperspectral imaging of tissue perfusion and oxygenation in wounds: assessing the impact of a micro capillary dressing. Journal of wound care 27, 38–51, doi:10.12968/jowc.2018.27.1.38 (2018).

18 Grambow, E. et al. Hyperspectral imaging for monitoring of perfusion failure upon microvascular anastomosis in the rat hind limb. Microvascular research 116, 64–70, doi:10.1016/j.mvr.2017.10.005 (2018).

19 Barberio, M. et al. HYPerspectral Enhanced Reality (HYPER): a physiology-based surgical guidance tool. Surgical Endoscopy 34, 1736–1744, doi:10.1007/s00464-019-06959-9 (2020).

20 Felli, E., Urade, T., Barberio, M., Felli, E. & Diana, M. Hyperspectral imaging of pig liver ischemia: a proof of concept, <https://www.airitilibrary.com/Publication/alDetailedMesh?docid=15610497-201912-201912180004-201912180004-117-121> (2019).

21 Holmer, A. et al. Oxygenation and perfusion monitoring with a hyperspectral camera system for chemical based tissue analysis of skin and organs. Physiological measurement 37, 2064–2078, doi:10.1088/0967-3334/37/11/2064 (2016).

22 Markgraf, W., Feistel, P., Thiele, C. & Malberg, H. Algorithms for mapping kidney tissue oxygenation during normothermic machine perfusion using hyperspectral imaging. Biomedical Engineering / Biomedizinische Technik 63, 557, doi:https://doi.org/10.1515/bmt-2017-0216 (2018).

23 Barberio, M. et al. Hyperspectral based discrimination of thyroid and parathyroid during surgery. 4, 399, doi:https://doi.org/10.1515/cdbme-2018-0095 (2018).

24 Jansen-Winkeln, B. et al. Determination of the transection margin during colorectal resection with hyperspectral imaging (HSI). Int J Colorectal Dis 34, 731–739, doi:10.1007/s00384-019-03250-0 (2019).

25 Köhler, H. et al. Evaluation of hyperspectral imaging (HSI) for the measurement of ischemic conditioning effects of the gastric conduit during esophagectomy. Surgical Endoscopy 33, 3775–3782, doi:10.1007/s00464-019-06675-4 (2019).

26 Maktabi, M. et al. Tissue classification of oncologic esophageal resectates based on hyperspectral data. International journal of computer assisted radiology and surgery 14, 1651–1661, doi:10.1007/s11548-019-02016-x (2019).

27 René, T. et al. O129 CLASSIFICATION OF BARRETT’S CARCINOMA SPECIMENS BY HYPERSPECTRAL IMAGING (HSI). Diseases of the Esophagus 32, doi:10.1093/dote/doz092.129 (2019).

28 Kilkenny, C., Browne, W., Cuthill, I. C., Emerson, M. & Altman, D. G. Animal research: reporting in vivo experiments: the ARRIVE guidelines. British journal of pharmacology 160, 1577–1579, doi:10.1111/j.1476-5381.2010.00872.x (2010).

29 Zach, C., Pock, T. & Bischof, H. 214–223 (Springer Berlin Heidelberg).

30 Wedel, A., Pock, T., Zach, C., Bischof, H. & Cremers, D. 23–45 (Springer Berlin Heidelberg).

31 Sánchez Pérez, J. a. M.-L., Enric and Facciolo, Gabriele. TV-L1 Optical Flow Estimation. Image Processing On Line 3, 137–150 (2013).

32 Chen, C. The art of bowel anastomosis. Scand J Surg 101, 238–240, doi:10.1177/145749691210100403 (2012).

33 Nickel, F. et al. Minimally Invasive Versus open AbdominoThoracic Esophagectomy for esophageal carcinoma (MIVATE) - study protocol for a randomized controlled trial DRKS00016773. Trials 22, 41, doi:10.1186/s13063-020-04966-z (2021).

34 Kesler, K. A. et al. Outcomes of a novel intrathoracic esophagogastric anastomotic technique. J Thorac Cardiovasc Surg 156, 1739–1745 e1731, doi:10.1016/j.jtcvs.2018.05.088 (2018).

35 Edholm, D. & Sundbom, M. Comparison between circular-and linear-stapled gastrojejunostomy in laparoscopic Roux-en-Y gastric bypass--a cohort from the Scandinavian Obesity Registry. Surg Obes Relat Dis 11, 1233–1236, doi:10.1016/j.soard.2015.03.010 (2015).

36 Radrich, K. & Ntziachristos, V. Quantitative multi-spectral oxygen saturation measurements independent of tissue optical properties. Journal of Biophotonics 9, 83–99, doi:https://doi.org/10.1002/jbio.201400092 (2016).

37 Eisel, M. et al. Investigation of optical properties of dissected and homogenized biological tissue. Journal of biomedical optics 23, 091418 (2018).

38 Liu, P., Zhu, Z., Zeng, C.-C. & Nie, G. Specific absorption spectra of hemoglobin at different PO2 levels: potential noninvasive method to detect PO2 in tissues. Journal of biomedical optics 17, 125002 (2012).

39 Eishi, H., Yamaguchi, K., Hiramatsu, Y. & Akita, K. Intra-mural distribution of the blood vessels in the stomach demonstrated by contrast medium injection: a cadaver study. Surg Radiol Anat 43, 389–396, doi:10.1007/s00276-020-02613-5 (2021).

40 Tomioka, K. et al. Morphometric and quantitative evaluation of the gastroepiploic artery. Okajimas Folia Anat Jpn 92, 33–35, doi:10.2535/ofaj.92.33 (2015).

41 Tomioka, K. et al. Anatomical and surgical evaluation of gastroepiploic artery. Okajimas Folia Anat Jpn 92, 49–52, doi:10.2535/ofaj.92.49 (2016).

42 Buunen, M. et al. Vascular anatomy of the stomach related to gastric tube construction. Diseases of the esophagus : official journal of the International Society for Diseases of the Esophagus 21, 272–274, doi:10.1111/j.1442-2050.2007.00771.x (2008).

43 Takeda, F. R., Cecconello, I., Szachnowicz, S., Tacconi, M. R. & Gama-Rodrigues, J. Anatomic study of gastric vascularization and its relationship to cervical gastroplasty. J Gastrointest Surg 9, 132–137, doi:10.1016/j.gassur.2004.03.006 (2005).

44 Kukar, M. et al. Minimally Invasive Ivor Lewis Esophagectomy with Linear Stapled Anastomosis Associated with Low Leak and Stricture Rates. J Gastrointest Surg 24, 1729–1735, doi:10.1007/s11605-019-04320-y (2020).

45 Okabe, H. et al. A long-term follow-up study of minimally invasive Ivor Lewis esophagectomy with linear stapled anastomosis. Surg Endosc, doi:10.1007/s00464-021-08482-2 (2021).

46 Barberio, M. et al. Quantitative serosal and mucosal optical imaging perfusion assessment in gastric conduits for esophageal surgery: an experimental study in enhanced reality. Surg Endosc, doi:10.1007/s00464-020-08077-3 (2020).

