## Supplement for "Optimization of anastomotic technique and gastric conduit perfusion with hyperspectral imaging in an experimental model for minimally invasive esophagectomy"

### Supplementary Text 1: Image Registration

When comparing hyperspectral recordings of the gastric conduit before and after application of the magnet, inevitably the contour slightly varied due to iatrogenic manipulation (**Supplementary Figure 1a and b**). In order to ensure an unbiased selection of the ROI on the control recordings of the gastric conduit, an image registration of both recordings was advised. To aid registration by algorithm, the contour of all gastric conduit recordings was annotated (**Supplementary Figure 1c**) and a transformation executed on the control recording for the best possible dynamic fit onto the intervention recording with magnet. This ensures a highly accurate overlay and pixel-wise comparability of both recordings (**Supplementary Figure 1d and e**). The rectangular measuring ROI was then manually placed next to the magnet and automatically transferred to the corresponding region on the registered control image. Subsequently, the $StO_2$ values of the annotated pixels could be extracted, presented in a histogram (**Supplementary Figure 1g**) and used for further calculations.

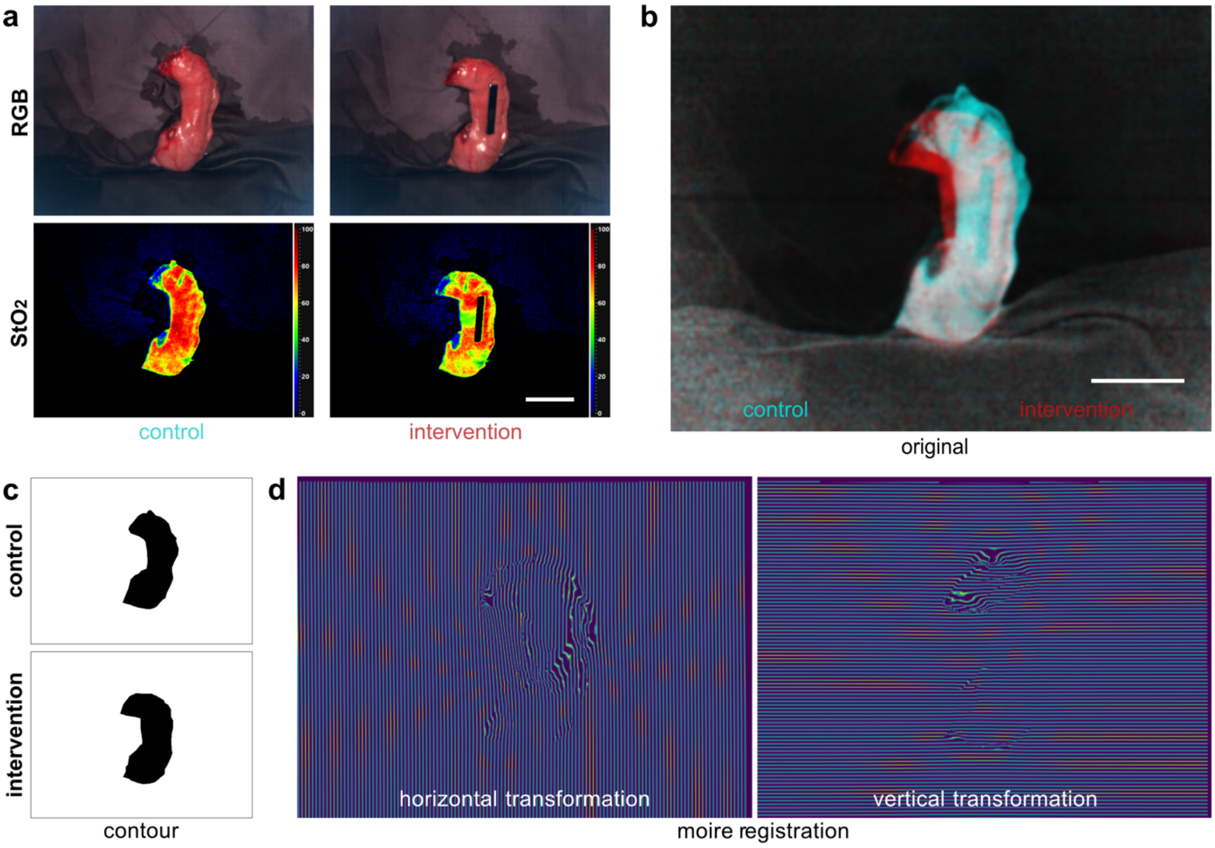

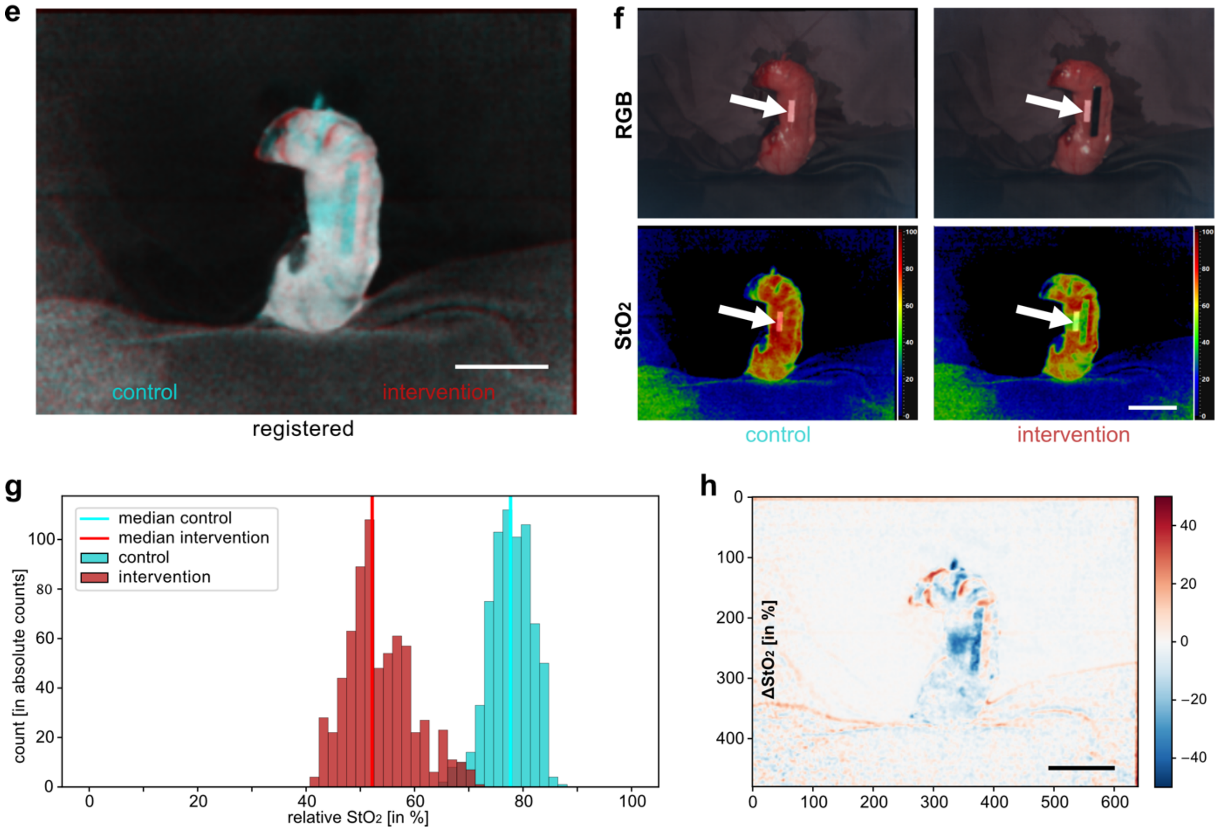


**Supplementary Figure 1 | Registration of hyperspectral recordings. a**, exemplary visualization of unregistered control and intervention**. b**, visualization of different contours**. c**, annotation of the conduit**. d**, moire registration with visualization of horizontal and vertical transformation**. e**, visualization of congruent contours. **f**, unregistered control and intervention with ROIs (arrows). **g**, histogram for $StO_2$ values of both ROIs. **h**, $\Delta StO_2$ image as numerical difference between control $StO_2$ and intervention $StO_2$.

**Supplementary Text 2: Model validation and additional phenomenon of gastroepiploic perfusion**

In order to validate the magnet-induced ischemia model as a proper model for stapler-induced ischemia of the gastric conduit during linear stapled anastomosis in Ivor-Lewis esophagectomy, there were two essential steps. The first step was the choice of the magnet. The strong neodymium magnet with the same dimensions as a linear stapler could ensure impairment of capillary flow by exceeding necessary capillary perfusion pressure by many times. In a second step, the response in measured $StO_2$ values was compared and could be identified as identical.

Subsequently to the median dissection at the usual position of the magnet, the incision was extended downward (**Supplementary Figure 2**) and oxygenation levels recorded. As expected, HSI oxygenation values dropped on the left side of the incision approximately up to the same transverse height as the incision (region I: 31.1±2.1%) while they remained high on the counterside (region II: 64.5±7.7% ($p=0.0068$) and region IV: 66.4±4.4% ($p=0.0018$)). However, repeatedly in the lower area of the gastric conduit, oxygenation values remained similarly high (region III: 63.9±8.2% ($p=0.0068$)). While stated p-values indicate a statistically significant difference between region I and the other regions, there were no significant differences within these other regions.

Although having only minor clinical relevance, this finding underlines the influence of caudal conduit perfusion via the right gastric artery and enhances the understanding of gastric conduit physiology.

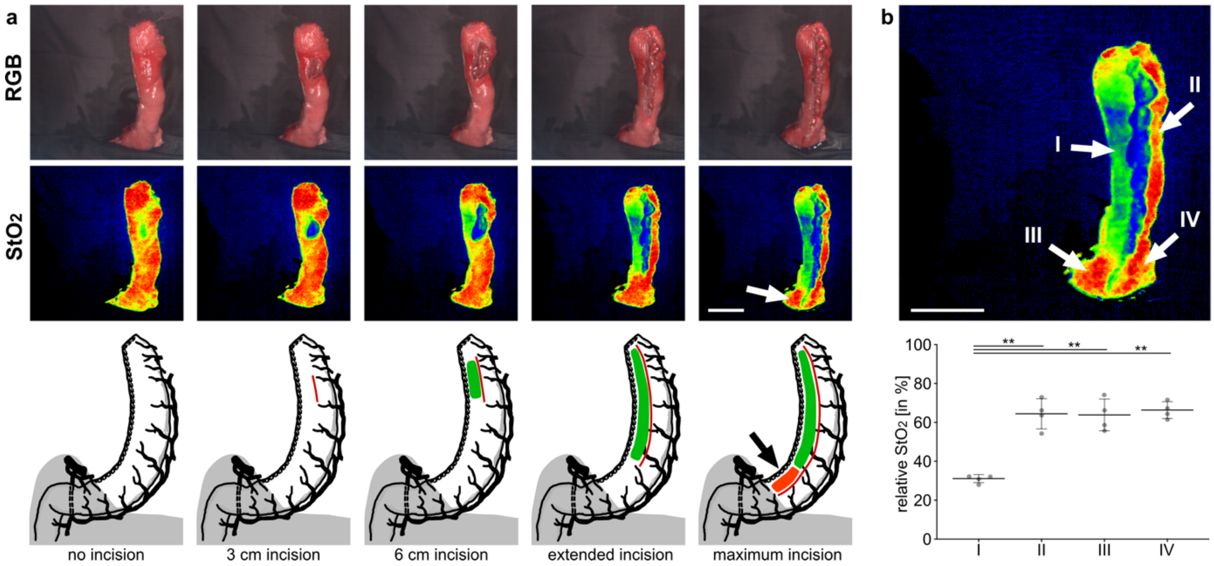


**Supplementary Figure 2 | Model validation and evaluation of the gastric conduit after median dissection. a,** evaluation of oxygenation of the gastric conduit after median dissection along its entire length (A=4; n=4)**. b,** Quantification of $StO_2$ levels of all four indicated ROIs. An ordinary one-way ANOVA was performed; ** is p≤0.01. Numbers in the box plots were obtained from the ROIs. Graphs depicts mean and standard deviation.

**Supplementary Text 3: Width of the gastric conduit did not influence tissue oxygenation**

Only short (3 cm) stapler simulations revealed significant differences in $StO_2$ values for cranial and caudal positions (**Figure 4 I and II**). Therefore, in order to evaluate further influencing factors for oxygenation differences within these two relevant groups, the results of these were stratified for gastric conduit width in a subgroup analysis (**Supplementary Figure 3**). Differences between control and intervention were still significant for the cranial position in thin conduits (inner diameter of 2 cm) with 63.1±5.5% vs. 47.5±6.2% (p=0.0031) (**Supplementary Figure 3 I**) as well as in wide conduits (inner diameter of 4 cm) with 69.4±7.1% vs. 53.8±8.6% (p=0.0305) (**Supplementary Figure 3 III**). Moreover, differences between control and intervention were still not significant for neither the caudal position in thin conduits with 68.5±6.8% vs. 68.0±10.3% (n.s.) (**Supplementary Figure 3 II**) nor in wide conduits with 74.4±3.5% vs. 74.1±4.4% (n.s.) (**Supplementary Figure 3 IV**).

Graphical baseline subtraction was performed (**Supplementary Figure 3d**) and groups were not significantly different after stratification for width i.e. thin cranial with -15.6±8.6% (**Figure Supplementary 3g.I**) compared to wide cranial with -15.6±14.6% (**Supplementary Figure 3g.III**) or thin caudal with -0.4.±5.5% (**Supplementary Figure 3g.II**) compared to wide caudal with -0.2±1.0% (**Supplementary Figure 3g.IV**). Analysis of the $StO_2$ levels of the whole unmanipulated conduit also revealed no significant difference between thin conduits (2 cm) (66.3±3.3%) and wide conduits (4 cm) (67.7±2.0%) (**Supplementary Figure 3h**). A detailed analysis of relative $StO_2$ values (**Supplementary**

**Figure 3i**) and ΔStO2 values of the ROIs (**Supplementary Figure 3j**) revealed no statistically significant difference between these two widths of gastric conduits.

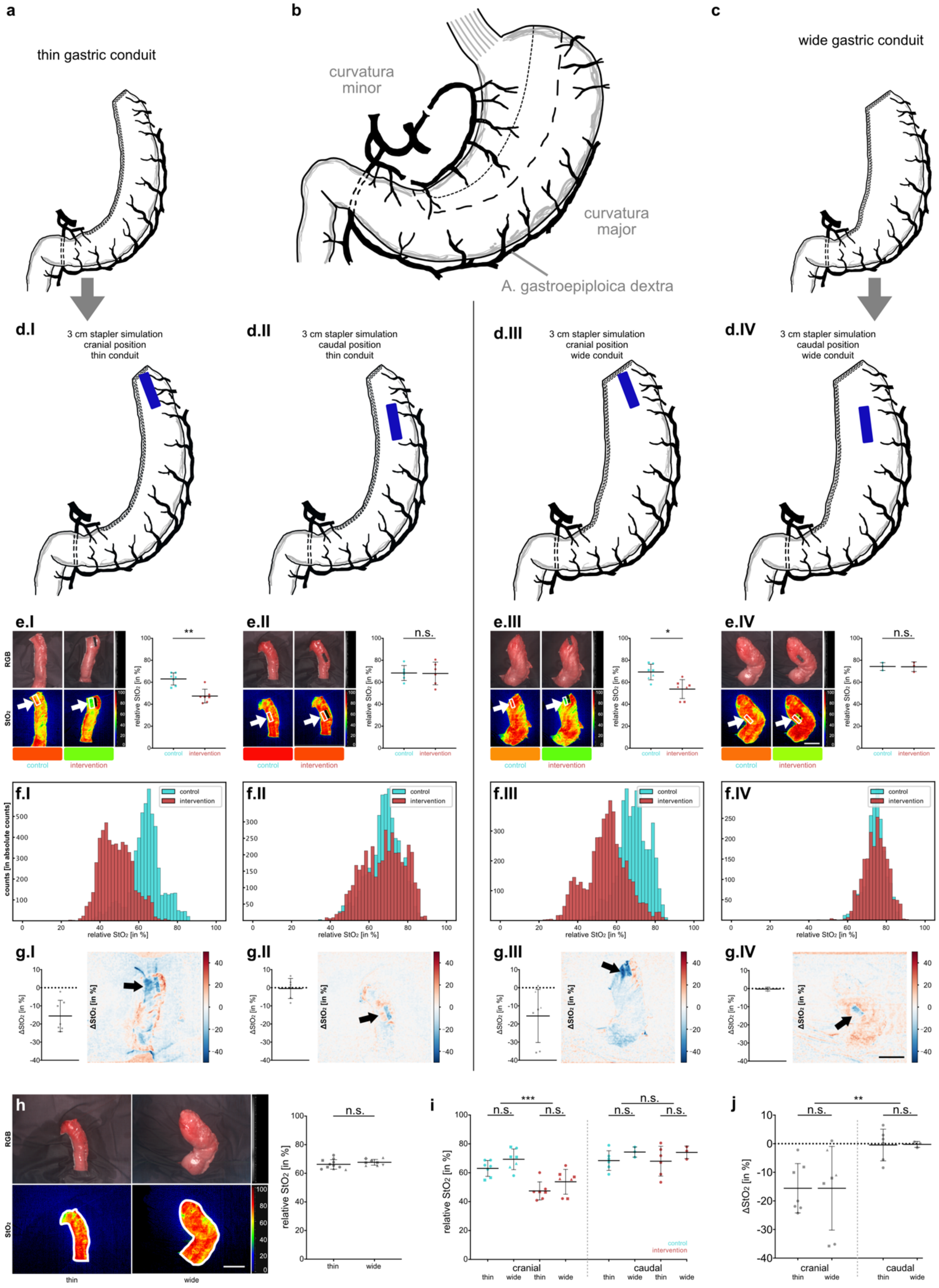


**Supplementary Figure 3 | Subgroup analysis for thin and wide gastric conduits.** Thin conduits had an inner diameter of 2 cm, while wide conduits had an inner diameter of 4 cm. **I**, thin short (3 cm)

cranial stapler simulation (A=6; n=7). **II**, thin short (3 cm) caudal stapler simulation (A=6; n=6)**. III**, wide short (3 cm) cranial stapler simulation (A=4; n=7). **IV**, wide short (3 cm) caudal stapler simulation (A=3; n=3). **a**, schematic drawing of a thin gastric conduit. **b**, schematic drawing of porcine stomach. **c**, schematic drawing of a wide gastric conduit. **d**, schematic drawing of linear stapling techniques. **e**, exemplary visualization of oxygenation and corresponding quantification. White boxes and arrows indicate ROIs for measurements. For **e** a paired t-test was performed. **f**, aggregated histograms of $StO_2$ values. **g**, quantification of the $StO_2$ difference in the ROIs between control and intervention and color-coded visualization. Black arrows indicate ROI next to the magnet. **h**, oxygenation of the whole unmanioulated gastric conduit comparing thin (A=8; n=11) and wide (A=4; n=9) indicating that there is no baseline difference between thin and wide. For **h** a paired t-test was performed. **i,** subgroup analysis of the quantification of **e**. **j,** subgroup analysis of the quantification of **g**. For **i** and **j** an unpaired t-test was performed; n.s. is not significant, * is p≤0.05, ** is p≤0.01 and *** is p≤0.001. Numbers in the box plots were obtained from the ROIs. Different animals are coded in different shapes; circles are different animals. Graphs depicts mean and standard deviation.

Therefore, this subgroup analysis between thin and wide conduit led to the conclusion that no difference in perfusion and anastomotic behavior could be seen. There was neither a relevant difference in HSI tissue oxygenation of the whole conduit prior to the stapling simulation (**Supplementary Figure 3h**) nor a relevant difference of the critical region on the conduit after the stapling simulation (**Supplementary Figure 3a-g and 3i-j**). However, this statement can only be made for inner diameters between 2 and 4 cm and not necessarily for conduits exceeding these diameters. Other retrospective studies found that wider conduits were prone to having a higher incidence of anastomotic insufficiency (17.3% vs. 8.7, p=0.041) [1], and that greater width of the gastric conduit was a significant risk factor for the development of anastomotic stricture (odds ratio = 3.36, *p*=0.005) [2]. Consequently, equal levels of HSI tissue oxygenation can only be seen as one of several contributing factors and besides equal levels of tissue oxygenation, thin gastric conduit construction seems clinically beneficial. Having identified the gastroepiploic artery as the main source of perfusion, it appears logical to construct a thin gastric conduit in order to decrease the length of necessary capillary perfusion along the transverse axis. This will have to be confirmed in clinical studies that will have to take into account functional outcome of the gastric conduit as well.

**Supplementary Text 4: Anatomical correlation to hyperspectral findings with vascular corrosion casting and scanning electron microscopy**

Evaluation of hyperspectral findings regarding tissue oxygenation suggests that the blood supply mainly comes from the gastroepiploic artery and travels transversely through the gastric conduit as opposed to travelling longitudinally from former gastric antrum to gastric fundus. Therefore, the question arose, whether an anatomical correlation providing a pathomechanistic explanation to these results could be found. As a result, vascular corrosion casting of the stomach was obtained using Biodur E20® (Biodur Products, Heidelberg, Germany). After euthanasia with i.v. potassium chloride, the abdominal aorta was cannulated, the distal thoracic aorta and iliac arteries were clamped and the inferior caval vein was incised. Perfusion was initiated by pressured infusion of 1,000 ml of

Sterofundin® (B. Braun®) through previously mentioned cannulation including 50,000 I.U. heparin, followed by 400 ml Sterofundin®. Biodur E20® for injection was mixed at a ratio of 100:45 (v/v) Biodur E20® Plus and catalyst E20 and injected by manual pressure. The infusion was stopped after venous return of the casting agent was observed in the inferior caval vein and the material initially hardened for several minutes without further manipulation. The gastric specimen was explanted and incubated for 12 h in a 40°C water bath for further hardening. Gastric tissue was removed with 15% (w/v) potassium hydroxide (RT; 3 days) and the resulting vascular corrosion cast was subsequently rinsed in water (**Supplementary Figure 4a-b**). SEM samples were prepared from corpus areas of stomach corrosion cast specimens. 20 mm x 10 mm samples of the outer layer were 10 nm gold/platinum (80:20) sputtered (Leica EM ACE 600, Leica Microsystems GmbH, Wetzlar, Germany) and analyzed by SEM (Zeiss Leo Gemini 1530, Carl Zeiss AG, Oberkochen, Germany). SEM images were taken at different magnifications with an accelerating voltage of 2.0 kV (**Supplementary Figure 4d-e**).

After corrosion casting of porcine stomach and SEM of cast gastric capillaries, it could be quantified that a significantly greater share of all vascular pixels (**Supplementary Figure 4c-d**) had a greater length of interrupted vascular neighbor pixels in the transversal orientation (67.4±10.5%) than in the longitudinal orientation (32.6±10.5%) ($p<0.0001$). This not only supports the hypothesis of predominantly transgastroepiploic blood supply of the conduit, but it also delivers pathomechanistic explanations to the observed phenomenon.

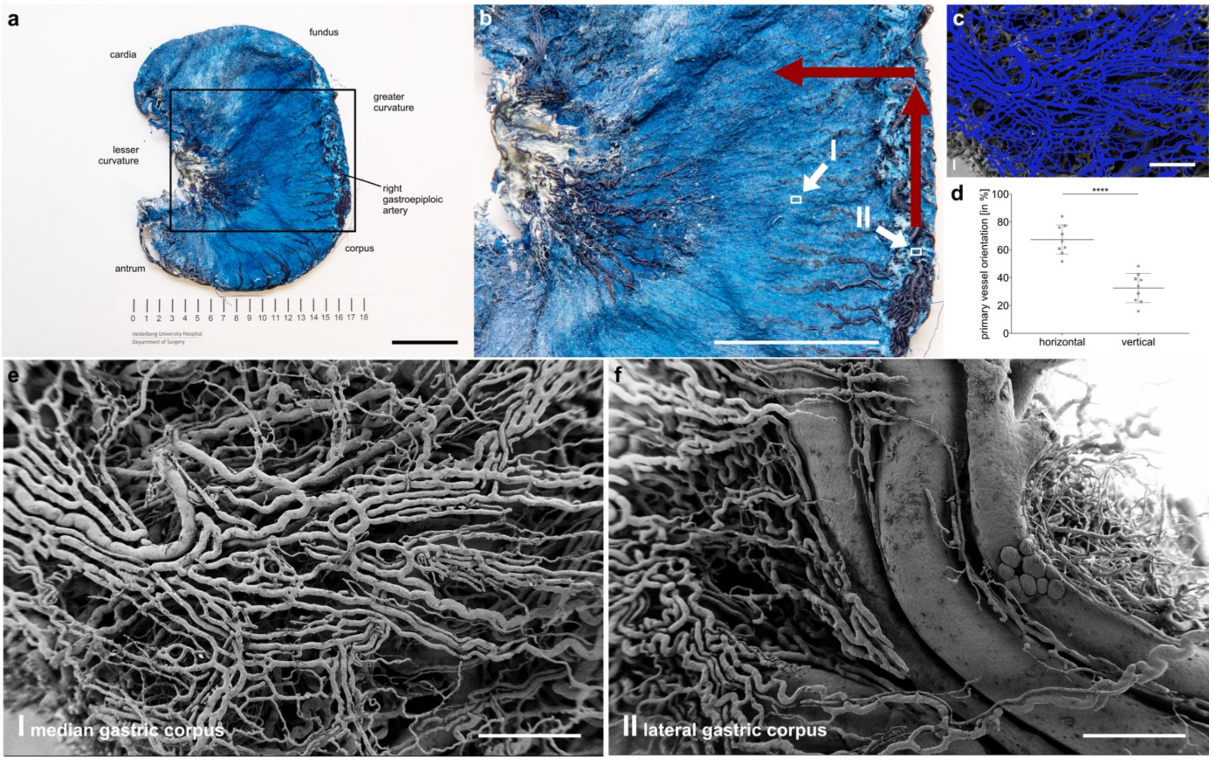


**Supplementary Figure 4 | Vascular corrosion casting of porcine stomach and scanning electron microscopy. a**, cast porcine stomach. **b**, magnification with assumed blood flow via gastroepiploic artery as indicated by red arrows. **c**, manual annotation of vessels from the median gastric corpus wall. **d**, quantification of vessel orientation separated in primarily transverse (from lateral to medial gastric edge) and longitudinal (from fundal to antral gastric edge) orientation depending on the longest distance of uninterrupted pixels in the annotation (A=2; n=9); different animals are coded in different shapes. **e**, SEM sample from median gastric corpus. **f**, SEM sample from lateral gastric corpus with visible gastroepiploic artery. Graph depicts mean and standard deviation. Unpaired t-test was performed; **** is $p \leq 0.0001$. Scale bar in microscopic images equals 1 mm.

**Supplement References**

1 Shen, Y., Wang, H., Feng, M., Tan, L. & Wang, Q. The effect of narrowed gastric conduits on anastomotic leakage following minimally invasive oesophagectomy. *Interact Cardiovasc Thorac Surg* **19**, 263-268, doi:10.1093/icvts/ivu151 (2014).

2 Zhu, D. S. *et al.* Wide Gastric Conduit Increases the Risk of Benign Anastomotic Stricture After Esophagectomy. *Am Surg* **86**, 621-627, doi:10.1177/0003134820923317 (2020).
